# Low Shear in Short-Term Impacts Endothelial Cell Traction and Alignment in Long-Term

**DOI:** 10.1101/2023.09.20.558732

**Authors:** Mohanish K. Chandurkar, Nikhil Mittal, Shaina P. Royer-Weeden, Steven D. Lehmann, Yeonwoo Rho, Sangyoon J. Han

## Abstract

Within the vascular system, endothelial cells (ECs) are exposed to fluid shear stress (FSS), a mechanical force exerted by blood flow that is critical for regulating cellular tension and maintaining vascular homeostasis. The way ECs react to FSS varies significantly; while high, laminar FSS supports vasodilation and suppresses inflammation, low or disturbed FSS can lead to endothelial dysfunction and increase the risk of cardiovascular diseases. Yet, the adaptation of ECs to dynamically varying FSS remains poorly understood. This study focuses on the dynamic responses of ECs to brief periods of low FSS, examining its impact on endothelial traction—a measure of cellular tension that plays a crucial role in how endothelial cells respond to mechanical stimuli. By integrating traction force microscopy (TFM) with a custom-built flow chamber, we analyzed how human umbilical vein endothelial cells (HUVECs) adjust their traction in response to shifts from low to high shear stress. We discovered that initial exposure to low FSS prompts a marked increase in traction force, which continues to rise over 10 hours before slowly decreasing. In contrast, immediate exposure to high FSS causes a quick spike in traction followed by a swift reduction, revealing distinct patterns of traction behavior under different shear conditions. Importantly, the direction of traction forces and the resulting cellular alignment under these conditions indicate that the initial shear experience dictates long-term endothelial behavior. Our findings shed light on the critical influence of short-lived low-shear stress experiences in shaping endothelial function, indicating that early exposure to low FSS results in enduring changes in endothelial contractility and alignment, with significant consequences for vascular health and the development of cardiovascular diseases.

## Introduction

Endothelial cells (ECs), which line the interior surface of blood vessels, are constantly exposed to the mechanical forces generated by flowing blood^1^. Among these mechanical stimuli, fluid shear stress (FSS) stands out as a pivotal factor^2^. FSS is responsible for initiating a cascade of intracellular signaling events that regulate various physiological processes, including vascular tone, inflammation, and angiogenesis^3-7^. Adequate FSS, typically within the range of 1-2 Pa and exhibiting a laminar flow pattern, triggers the release of vasodilators such as nitric oxide, which facilitates the precise adjustment of blood vessel dilation and sustains optimal blood flow^8^. Furthermore, it fosters the expression of anti-inflammatory factors while discouraging the adhesion of circulating cells to the endothelial surface, thereby mitigating the risk of vascular ailments, including atherosclerosis^9-11^. Conversely, improper FSS, characterized by low levels (< 0.5 Pa) and/or disturbed shear patterns, instigates endothelial dysfunction^10,12,13^. This disturbance in FSS equilibrium disrupts the delicate balance between pro-inflammatory and anti-inflammatory factors, resulting in cardiovascular disorders^8^. Mechanisms by which endothelial cells sense and respond to low FSS encompass endothelial surface molecules, ion channels^14^, mechanosensors like vascular endothelial cadherin^15^, and downstream signaling pathways, including Rho-ROCK^16^, NF-κB^17^, and Smad2/3 pathways^18^. Dysregulation of these signaling pathways due to sustained or repeated exposure to low FSS can compromise vascular barrier function and contribute to the initiation and progression of atherosclerosis^19,20^.

Considerable research efforts have been directed toward comprehending the individual impacts of high or low fluid shear stress (FSS) on endothelial function^8,15,16,18,21-27^. However, the consequences of transient exposure to low FSS have remained elusive. The flow dynamics within the vascular system exhibit a constant state of flux, evolving over time^28^. Transient episodes of low FSS occur in scenarios characterized by disrupted flow patterns or in regions where blood flow experiences temporary reductions, such as during vasoconstriction or occlusion^29^. Furthermore, it is commonly recommended that *in vitro* flow experiments employing a parallel plate chamber involve a pre-conditioning phase for endothelial cells (ECs) under low flow speed initially with a gradual increase in flow rate afterward, as opposed to an abrupt application of high FSS^30-32^. This approach is advocated to minimize the potential detachment of cells. However, it is noteworthy that numerous studies based on parallel-plate systems appear to directly apply designed laminar high FSS without reference to such pre-conditioning practices^27,33-36^. Yet, a comprehensive understanding of how ECs respond to temporally fluctuating FSS, commencing with low magnitudes, remains limited.

To investigate the tension state of the endothelial monolayer, our study employs a method called traction force microscopy (TFM)^37^. This technique allows for the measurement of traction exerted by ECs on an underlying soft substrate. The traction represents a functional outcome of the contractile status of the monolayer as well as cell adhesion to the extracellular matrix and neighboring ECs, which is governed by the FSS-induced signals such as NF-kB, YAP and RhoA, and eNOS activity^8,15,38-43^. Traction of ECs has been associated with a gap formation of an EC monolayer^44^. Nevertheless, traction response in response to high FSS has been in debate. Earlier studies suggested that both individual ECs and EC monolayers increase their traction in response to high FSS^25,27,40^. Subsequent investigations revealed that the traction of EC monolayers decreases under prolonged (>12 hour) exposure to high FSS^15,45^. Yet, the dynamics of traction over time, spanning from short to long-term exposure to high FSS, remain unclear. Furthermore, compared to high FSS, there has been a limited exploration of low FSS’s impact on traction. While one study indicated a pronounced increase in traction upon the introduction of low FSS, which was reversible with time^26^, whether this low FSS exerts a lasting effect on traction following a return to high FSS conditions has remained unexplored.

ECs exhibit a tendency to align themselves along the direction of long-term high FSS, whereas such alignment is notably absent in the presence of low FSS^21,23^. The alignment of ECs with the flow direction is closely linked to the downregulation of inflammatory pathways^46,47^ whereas misalignment of ECs triggers the activation of these inflammatory pathways^27,48,49^. Clinical observations have revealed that atherosclerotic regions in vivo are associated with the failure of endothelial cells to align and elongate in the direction of flow^50,51^. ECs aligned via micropatterning, after which are exposed to the flow in the aligned direction, have shown decreased inflammatory signaling whereas applying flow in the direction perpendicular to the alignment has promoted inflammatory signaling^52^. In static culture, ECs orient themselves randomly, but under the influence of low FSS, they exhibit alignment perpendicular to the flow, which is again linked to inflammatory responses^21,23^. Notably, a recent study employing TFM has provided indirect evidence suggesting that traction alignment may serve as a precursor to EC alignment with the flow direction^45^. However, the precise mechanisms governing traction alignments during the transition from low to high FSS have remained elusive.

In this study, to scrutinize EC traction at a finer temporal resolution during transitions from low to high shear stress, we integrated TFM with the custom-built flow chamber system. Our system allows for real-time monitoring of both short-term and long-term traction modulation of human umbilical vein endothelial cells (HUVECs) under time-varying FSS. Using this system, we find that HUVECs show a first transient rise in traction immediately upon exposure to low FSS. Prolonged observations unveiled that this traction continued to rise for over 10 hours, gradually decreasing but never returning to the baseline level even after 24 hours. In stark contrast, HUVECs exposed to high FSS exhibit a rapid rise first but also a rapid decline in traction within the first hour, then drops well below the no-shear stress baseline within 2 hours. We show that this behavior of traction magnitude is associated with the distribution of traction orientation: HUVECs previously subjected to low FSS retained traction orientations perpendicular to the flow direction, even after being subjected to high FSS for more than 20 hours. In contrast, cells initially subjected to high FSS aligned their traction forces with the flow direction, maintaining this alignment over both short and long durations. Through deep-learning-based cell segmentation analysis, we show high correlation of the traction alignment with the cell alignment. By adopting Granger Causality^53^, we show that traction alignment functionally causes cell alignment under direct high FSS and 60-min low FSS. Collectively, our findings suggest that initial exposure to low shear stress initiates a gradual modulation of traction, which takes a considerably longer time to return to a relaxed contractile state under high shear stress conditions and traction is a precursor for cell alignment.

## Materials and methods

### Flow system with traction microscopy

A rectangular flow chamber was designed to have 38 × 5 × 0.8 mm of inner flow channel dimensions and was manufactured using polycarbonate via CNC machining. The channel dimension allows for effective imaging by enabling to position the viewing area closer to the microscope objective. Overall dimensions of the flow chamber were considered based on the confined dimensions of the stage top incubator with a height of 25 mm and an insert of 75 x 25 mm glass slide. The outer dimension of the flow chamber has a height of 10 mm and a rectangular dimension of 75 x 25 mm. This flow chamber was attached to the bottom coverslip via adhesive tape (3M).

Before attachment to the flow chamber top, the cover glass was prepared to contain a silicone gel with fluorescent markers for traction measurement. Coating with silicone gel and fluorescent beads was done as previously done^54^. Briefly, Q-gel 920A and 920B (Quantum Silicones LLC, Richmond VA) were mixed in 1:1.2 proportion which yields 2.6 kPa of shear modulus. 300 mg of the mixture was added to the coverslip and spin-coated at an acceleration of 1000 rpm and rotation of 2500 rpm for a uniform thickness of 20-25 µm. The coverslip was then cured at 80℃ for 2 hours. After curing, the substrate was treated with (3-Aminopropyl) triethoxysilane (Sigma-Aldrich), diluted at 5% in absolute ethanol, and then treated for 3 minutes. The substrate coverslip was then rinsed with 96% ethanol 3 times and air dried for 10-15 mins. For bead coating, 5000 µL of sterile DI (di-ionized) water was added with 2.5 µL of 200 nm-diameter dark-red (with 660 nm excitation and 680 nm emission wavelengths) microspheres beads (FluoSpheres™) and sonicated for 15 mins. The observed density of beads was around 2 beads/µm^2^. After sonication, the dried coverslip substrate was treated with the bead solution for 10 mins and rinsed 3 times in DI water.

The assembled flow chamber was integrated into the flow system that consists of a peristaltic pump Model# NE-9000, New Era Pump Systems, Inc., USA), a pulse dampener, and a reservoir (Fig. 1a). The peristaltic pump was used because it supports a wide range of flow rates and allows minimal backpressure during the uniform media circulation. The pulse dampener was used to eliminate the pulsation in the flow due to the pump as well as to ensure a continuous flow of cell media without any air bubbles. A glass bottle was used as a reservoir, which has three holes for the inlet, outlet, and CO_2_. The CO_2_ concentration and the temperature inside the chamber were maintained by enclosing the media inside the CO_2_ incubator. The pump was operated using LabVIEW for flow profile programming flexibility. The flow system was maintained at 37 °C and 5% CO_2_ throughout experiments (Fig. 1a).

**Figure 1.**
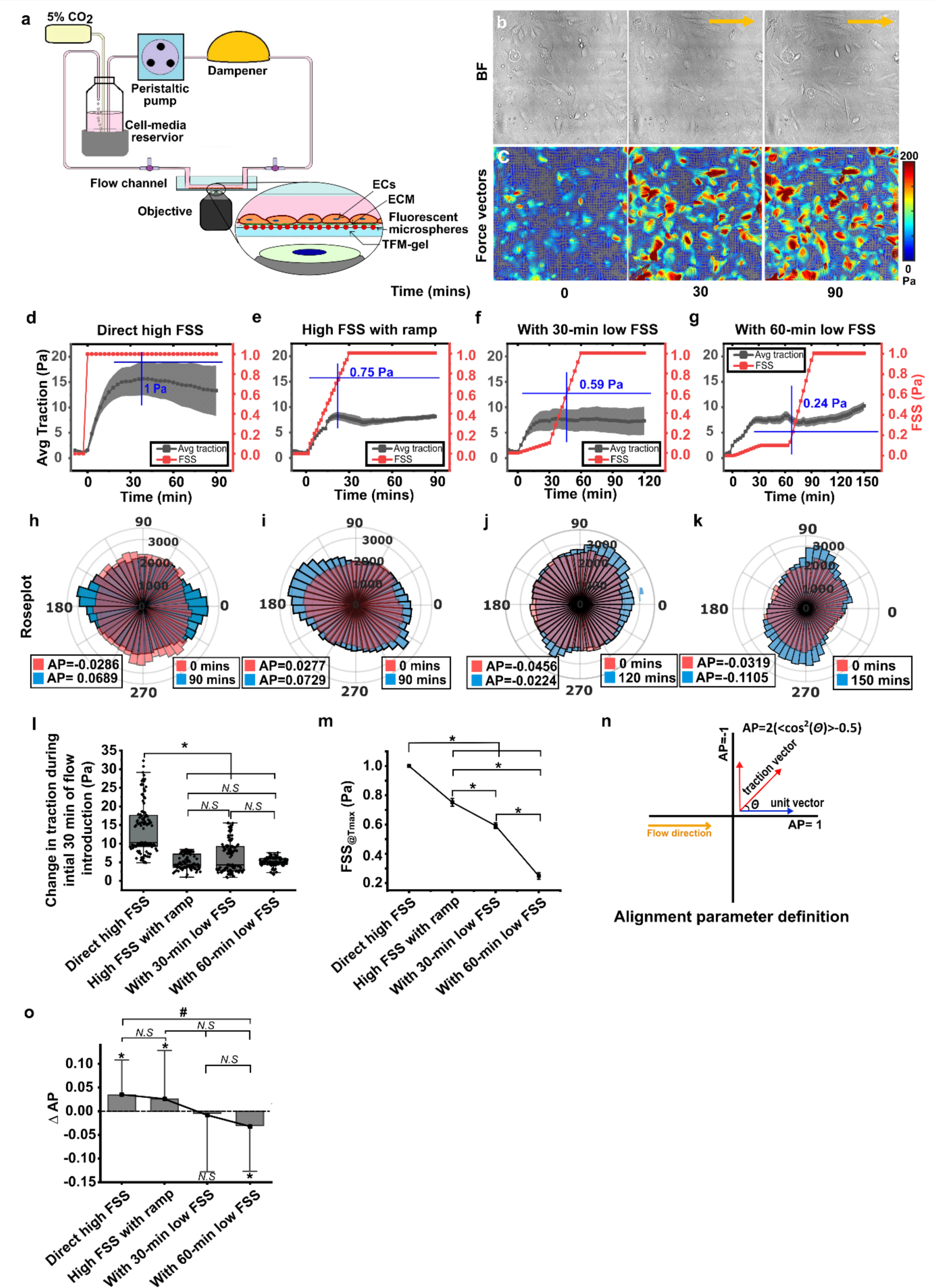
Low FSS increases traction magnitude but induces the orientation of tractions to be perpendicular to the flow in HUVECs. (a) A closed-loop flow system consisting of a programmable peristaltic pump, a flow dampener, a flow channel, and a media reservoir. A zoomed-in view highlights the EC monolayer on bead-coated gel in the custom-built flow channel. (b, c) DIC images (b) and color-coded traction vector fields (c) at 0 mins (before the onset of the flow), 30 mins, and 90 mins of a region of interest of a HUVEC monolayer after direct high FSS flow application. A yellow arrow indicates the flow direction. (d-g) Average traction magnitude, black, as a function of time, co-plotted with FSS profiles on ECs in red for direct 1 Pa FSS (d, M=1, N=6), 1 Pa FSS with 30 min ramp (e, M=1, N=6), with 0.1 Pa FSS 30-min ramp (f, M=1, N=6), and with 60-min 0.1 Pa shear (g, M=1, N=6). Error bars: S.E.M. (h-k) Polar histograms of traction fields before (red) and after (blue) the duration of the flow for each condition under d-g with alignment parameter (AP_traction_). (l) A boxplot of average traction changes during the first 30 minutes of initial flow introduction. M=3, N=16 for direct high FSS, M=3, N=20 for ramped high FSS, M=3, N=18 for a flow with 30-minute low FSS, M=3, N=16 for a flow with 60-minute low FSS. (m) A plot of FSS_@Tmax_ under all flow conditions. *: p < 0.05 tested with one-way ANOVA with Tukey’s post-hoc test. M, N are the same as in panel l. (n) Illustration about the definition of AP_traction_ of three traction vectors to the flow direction. One vector in the flow direction, another perpendicular to the flow direction, and one arbitrary direction with an angle 8. (o) A bar plot of changes in AP_traction_ before and after each flow condition. Error bars: S.E.M. *: p <0.05 tested with 1-sample t-test, *i.e*., compared against zero. #: p < 0.05 tested with one-way ANOVA with Tukey’s post-hoc test. M and N are the same as in panel l. M represents the number of flow experiments and N represents the number of regions of interest observed in each experiment.

### Cell culture and seeding to the flow chamber

The Human Umbilical Vein Endothelial Cells wild type, Pooled (HUVECs) cell line was obtained from Zhao Lab (MTU, MI) and was originally purchased from Lonza (cat#: C2519A). The cells were used from passages 5-9 for the experiments and were cultured in endothelial cell growth Medium-2 (EGM™-2) Bulletkit™ (Lonza Bioscience) on pretreated tissue culture 6-well plate (VWR international). For the flow experiments, 0.75×10^6^ cells/ml were seeded on the soft gel-coated coverslip and stored in a CO_2_ incubator for 12 hours to form a monolayer before imaging. Before cell seeding, the bead- and gel-coated cover glass was attached to the flow channel using biocompatible adhesive tape with a thickness of 5-7 µm. Afterward, to facilitate HUVEC adhesion, the silicone gel housed with the flow chamber was coated with 0.5 μg/mL of fibronectin (FN) via 1-ethyl-3-(3-dimethylaminopropyl) carbodiimide hydrochloride (EDC) chemistry as in bead coating process. Specifically, 10 µL of 10 mg/mL EDC solution was added to 1 mL PBS, to which 5 µL of 10 µg/mL fibronectin stock solution was added via flow inlet port. The FN-EDC mixture was incubated on the gel for 30 minutes at room temperature and rinsed once with PBS. To block unspecific binding and remove the excess FN, the substrate was incubated at room temperature for 30 mins with 1 mL of 0.1% BSA in PBS and rinsed 3 times in PBS.

### Shear flow experiment

For the shear flow experiments, the calculations for FSS were performed using the Poiseuille law modified for parallel plate flow channels^55^:

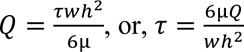

where Q is the desired flow rate, τ is the wall shear stress on the gel and cell surface, µ is the viscosity of the cell media (0.72 mPa·s), *w* is the width of the flow channel (5 mm), and h is the height of the channel (0.8 mm). Accordingly, to apply 1 Pa FSS, the flow rate of 44.4 ml/min was designed and applied. For 0.1 Pa FSS, 4.4 ml/min was applied and the ramp and low FSS profile were calculated and applied based on the time-dependent FSS increase from 0.1 Pa to 1 Pa. The flow was applied at least 30 minutes after placing the flow channel inside the incubation chamber to allow stable temperature and CO_2_ environment. The flow channel was supplied with respective FSS using a peristaltic pump operated using LabView software. Briefly, Laboratory Virtual Instrument Engineering Workbench (LabVIEW), a programming language designed for instrument control using graphical block diagrams connected using different function nodes via drawn wires. The user interface (front panel) is used to control each flow condition where in the background different flow profiles were designed. The front panel uses inputs that allow the user to supply information such as establishing the serial port communication between the pump and the system for automation of the process. The peristaltic pump provided commands as per the user manual were used to control the start and stop function of the pump. For the development and automation of each flow profile, the flow rates in ml/min were calculated based on the FSS calculations to achieve the desired FSS. The flow rates were embedded using the loop function and commands to increase or decrease the flow rates with respect to time. Thus, reducing the effort to manually control the pump flow rates and operation variability and errors.

### Live-cell imaging, traction reconstruction and analysis

The beads in the flow chamber were imaged via spinning-disk confocal microscope (Yokogawa CSU-X1, built on Nikon Ti-S) with 20x objective and 642 nm laser excitation before and after the flow application at every 180 seconds. At the same time, differential interference contrast (DIC) images were taken to observe the cell confluency, orientation, and alignment. The reflected light from SD Confocal is acquired by the ORCA-Flash sCMOS camera. Image acquisition was controlled by Metamorph software. During imaging, the flow chamber was kept inside a stage-top incubator (OKO lab) using a sealing putty to ensure stable imaging without drifting. For TFM imaging, after imaging in cell presence, HUVECs were removed from the substrate surface by flowing 0.25% trypsin into the chamber, and then the bead positions in relaxed gel configuration were imaged via far-red laser again.

The acquired images of the beads on the deformed gel (when HUVECs were present on the substrate) and of the reference configuration of the relaxed gel (after removing HUVECs) were processed for traction reconstruction described in Han et al., 2015^37^. In short, particle image velocimetry (PIV) was used to compare the bead images for deformed and relaxed gel. A template size of 17 pixels and a maximum displacement of 40 pixels was used based on the bead density and overall deformation. The subpixel correlation via image interpolation method was used for individual bead tracking for correction of bead tracking and removal of outliers before traction estimation. For bead images of challenging displacement field, we used our PTVR (particle tracking velocimetry with re-tracking) method to rescue potentially missed displacement vectors^56^. Boussiness equation assuming infinite gel thickness was used for solving the force field (defined as the expected deformation of the gel with all the bead locations applied with unit force at a particular force grid mesh). Further, the generated forward map was used to solve inverse problems based on the measured bead displacements to calculate the force field. The Regularized Fourier Transform Traction Cytometry (regFTTC) method was used as a reconstruction method. A detailed description of this method is present in Sabass et al., 2008^57^. Briefly, regFTTC solves the inverse problem by transforming it into the frequency domain. This method was used to reduce computation time and the priority was to quantify and compare the overall traction.

For visualization of the orientation of the traction vectors, the generated force field was used to calculate the angular distribution of the force alignment using a custom-written code in MATLAB. The vector alignment to the flow direction was calculated using the alignment parameter, AP_traction_ = 2(<cos^2^θ>-0.5), where θ represents the angle of a traction force vector to the flow direction, which is an x-direction in our configuration. The average of the distribution of angles for the force alignment was denoted using < >. AP_traction_ ranges between 1 to -1, where 1 represents a perfect parallel to flow alignment, -1 as perpendicular to flow alignment and 0 being isotropic alignment. This definition of AP was adapted from the definition of the orientation parameter. For traction rise comparison during the ramp high FSS and early low FSS condition, the change in traction within the first 30 mins were calculated. For calculation, the difference between the baseline traction before FSS and the traction between 24-30 mins after FSS were calculated to get the magnitude increase in each individual condition. The comparison of the traction rises with respect to zero were compared for significance. For the observed plateau phase during traction increase under rising FSS, FSS_@Tmax_ was calculated. For calculation, the difference between the traction with successive timepoint was compared for zero or negative to determine the maximum traction time point. Further, the comparison was made between the conditions to determine the FSS for each traction maximum based on the timepoint and applied FSS.

### Cell segmentation, training, and orientation analysis

The Cellpose 2.0^58^ was used with human-in-loop model to predict cell segmentation. Briefly, Cellpose 2.0 is a U-net architecture-based deep-neural network model with pretrained Cellpose 1.0 model utilization^59^. The training was done on the brightfield images of HUVECs for each subset of images and the prediction was further improved with manual training and correction. The models were trained from scratch for 100 epochs with batch size of 15, with weight decay of 0.0001 and a learning rate of 0.1. For all the training images with less than 5 regions of interest were excluded. For the analysis of the segmentation, the angular distribution of the orientation of all segmentations were calculated using a custom-developed code in python. The cell alignment to the flow direction was calculated using the alignment parameter, AP_cell_=2(<cos^2^θ>-0.5), where θ represents the angle of the segment’s major axis, after fitting an ellipse to each segmentation, to the flow direction, *i.e.*, the x-direction in our configuration. The polar color wheel with colormap was used to denote the angle of the predicted cell orientation for each condition. The comparison of AP_cell_ and AP_traction_ was made for each condition to determine the traction and cell alignment correlation. For AP_traction_, only prominent traction vectors were used, i.e., by subsampling with Otsu thresholding, to avoid noise-like effect from small traction vectors. The linear correlation between AP_traction_ and AP_cell_ was measured using Pearson correlation coefficient. The number between 1 and -1 showed the strength and direction of relationship between AP_traction_ and AP_cell_, where 1 represents strong positive correlation and -1 represents strong negative correlation.

### Statistical analysis

The data were obtained from 3 replicate experiments and 5 different positions in each experiment stated otherwise. Error bars in all figures represent the standard error of the mean and the analysis was done using OriginLab (Fig. 1d-g, Fig. 2a-c, Fig. 3a-d, Fig. 4j-m). The polar histogram was plotted to visually inspect the force-flow alignment and cell alignment using MATLAB function, ‘polarhistogram’ (Fig. 1h-k, Fig. 2d-f, Fig. 3e-h, Fig. 4f-i). To determine significant difference between groups, we used one-way ANOVA with Tukey’s post-hoc test (Fig. 1l, m, p, Fig. 2h, Fig. 3i, j, k, significance denoted by asterisk). To determine if the analyzed data *e.g.*, change in traction or change in AP are significantly more than zero, we used one-sample Student’s t-test per each group (Fig. 1o, Fig. 3k, significance denoted by #). P-values less than 0.05 were considered significant. The non-linear fit was done using Gaussian process regression (GPR) (Fig. 4j) in OriginLab. R-squared (R^2^) value was reported for non-linear curve fitting of the data. ANOVA with Tukey’s post-hoc test and Student’s t-test were done in JMP-pro software. Pearson correlation coefficient was used to report the correlation between AP_traction_ and AP_cell._. To determine if such correlations were significantly different from zero, we used a robust correlation test^60^, which accommodates possible dependence and heteroskedasticity in the two time-series. Within each experiment condition, we combined the correlation test results using the Harmonic Mean P (HMP)^61^ method. The HMP method is notable for its capability to combine multiple tests without requiring knowledge of the dependence structure between the tests. In each condition, AP_traction_ and AP_cell_ were to have a significant correlation if the HMP-combined p-value was less than 0.05 (Fig. 4n).

**Figure 2.**
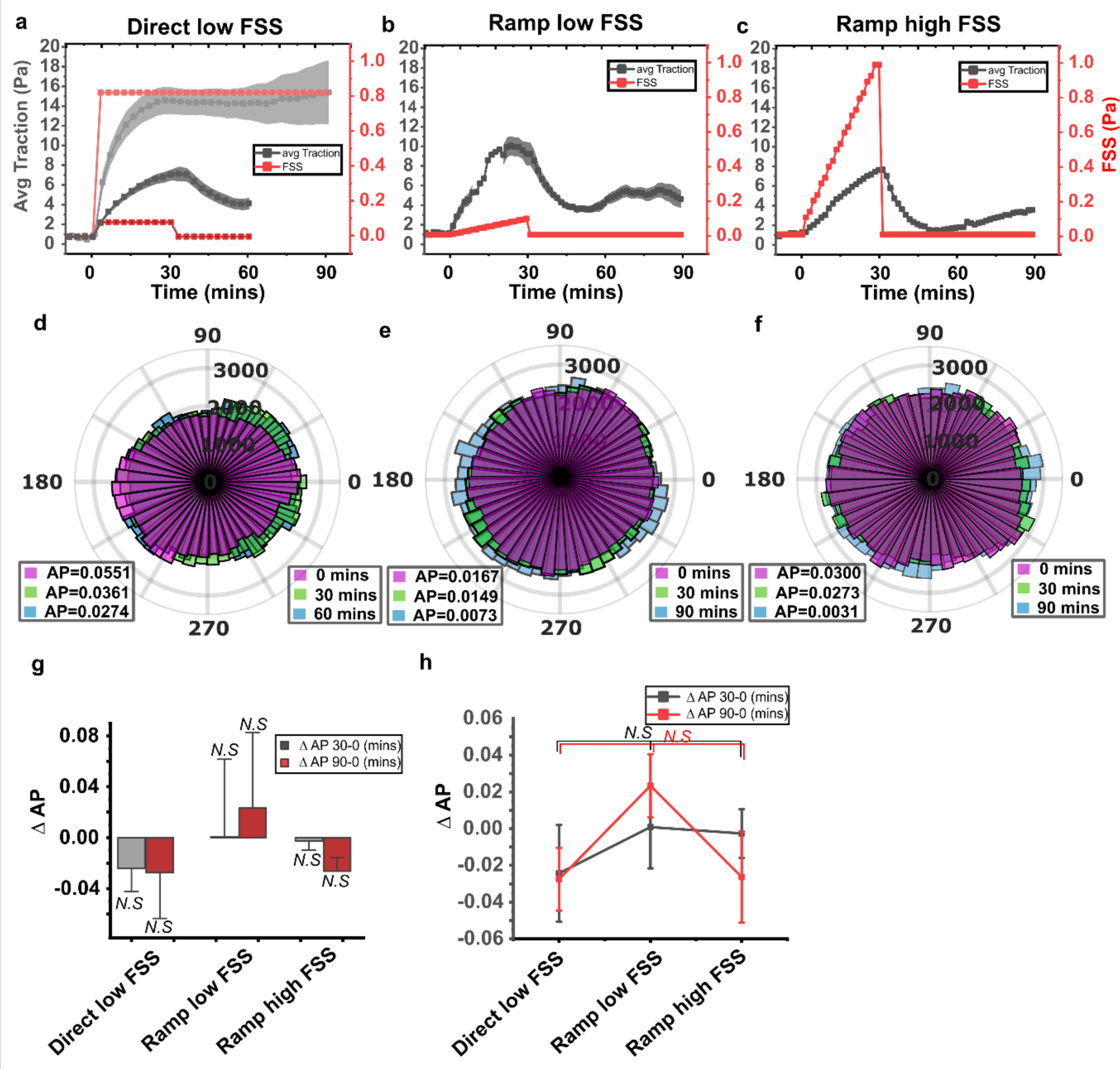
EC traction starts to decrease immediately after removal of low or high FSS. (a-c) Plots of the average traction magnitude, normalized by subtracting the baseline traction, over time under flows with 0.1 Pa FSS (a), ramped 0.1 Pa FSS (b), and ramped 1 Pa FSS (c). (d-f) Plots of polar distributions of traction field before and after flows with 0.1 Pa FSS (d), ramped 0.1 Pa FSS (e) and ramped 1 Pa FSS (f), recorded at 0 min (magenta), 30 min (light green), and 60 min (light blue). (g) A bar plot of changes in AP_traction_ after flow and after stopping the flow. *: p<0.05 tested with 1-sample t-test, compared against zero change. (h) A scatter plot of AP_traction_ for comparison between flow conditions after flow and after stopping the flow. *: p < 0.05 and *N.S* (not significant), tested by one-way ANOVA with Tukey’s post-hoc analysis. M=1, N=5 where, M represents the number of flow experiments and N represents the number of regions of interest observed in each experiment.

**Figure 3.**
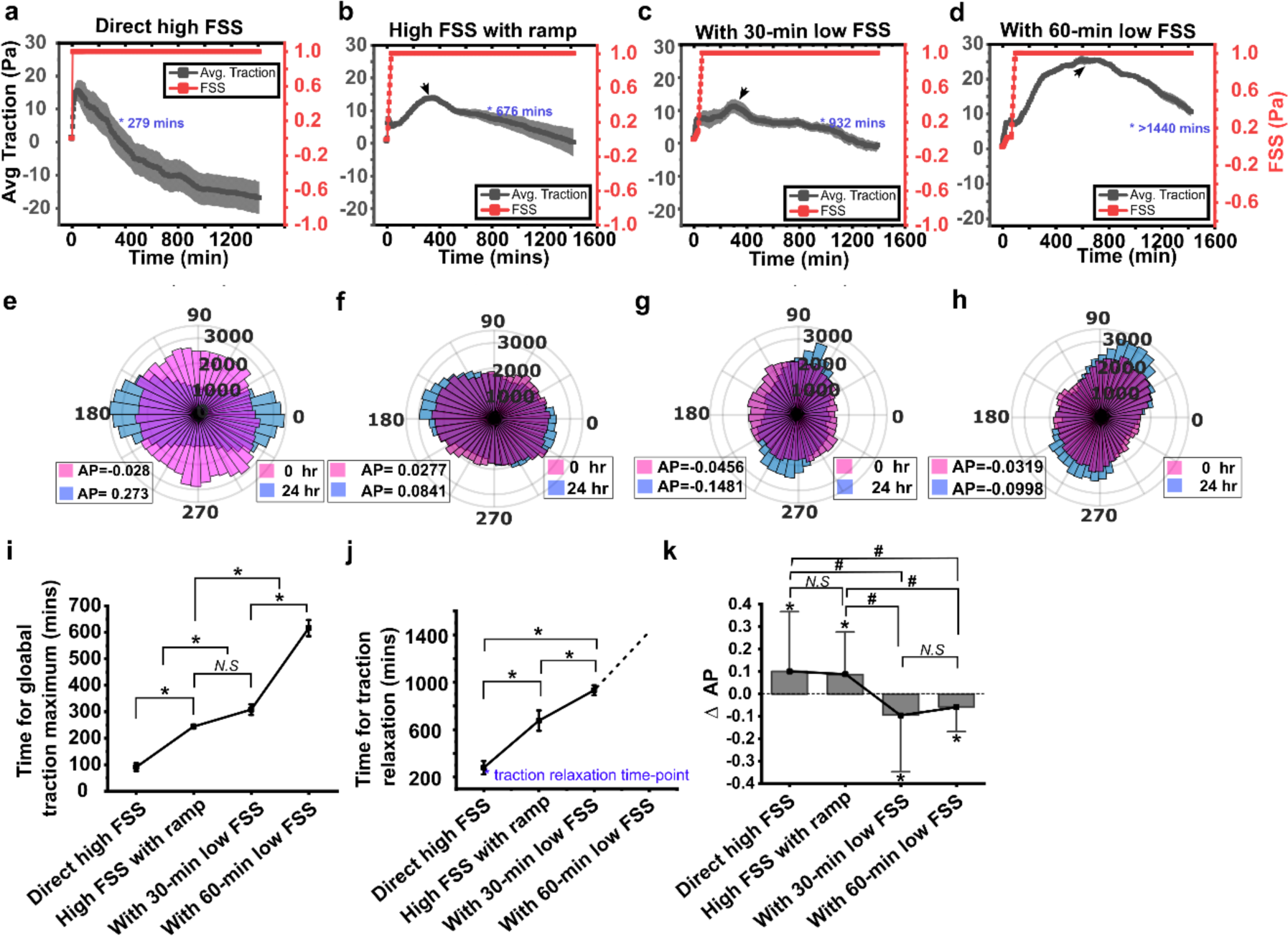
Exposure to early low FSS hampers ECs’ long-term traction relaxation and alignment to the flow under high FSS. (a-d) Average traction magnitude as a function of time, co-plotted with FSS profiles on ECs for direct 1 Pa FSS (a, N=6), 1 Pa FSS with 30 min ramp (b, N=6), with 0.1 Pa FSS 30-min ramp (c, N=6), and with 60-min 0.1 Pa shear (d, N=6). An arrowhead in each panel represents the time of global maximum. Blue asterisks and time information represent the traction relaxation time when the traction reached 110% of the baseline traction. Error bars: S.E.M. (e-h) Polar histograms of traction fields before (red) and after (blue) the duration of the flow for each condition under a-d with AP_traction_. (i) A scatter plot with mean ± s.e.m. showing time to achieve global traction maximum for each flow condition (direct high FSS (M=3, N=16), ramped high FSS (M=3, N=20), with 30-min ramped low FSS (M=3, N=18), and with 60-min low FSS (M=3, N=16). *: p < 0.05 tested with one-way ANOVA with Tukey’s post-hoc test. (j) A scatter plot with mean ± s.e.m. showing traction relaxation time when traction reduce to 110% of the baseline traction, also highlighted in (a-d) with *blue asterisk with time.* *: p < 0.05 tested with one-way ANOVA with Tukey’s post-hoc test. M, N are the same as in panel i (k) A bar plot of changes in AP_traction_ before and after each flow condition. *: p<0.05 tested with 1-sample t-test. Error bars: s.e.m. #: p < 0.05 tested with one-way ANOVA with Tukey’s post-hoc test. M and N are the same as in panel i. M represents the number of flow experiments and N represents the number of regions of interest observed in each experiment.

**Figure 4.**
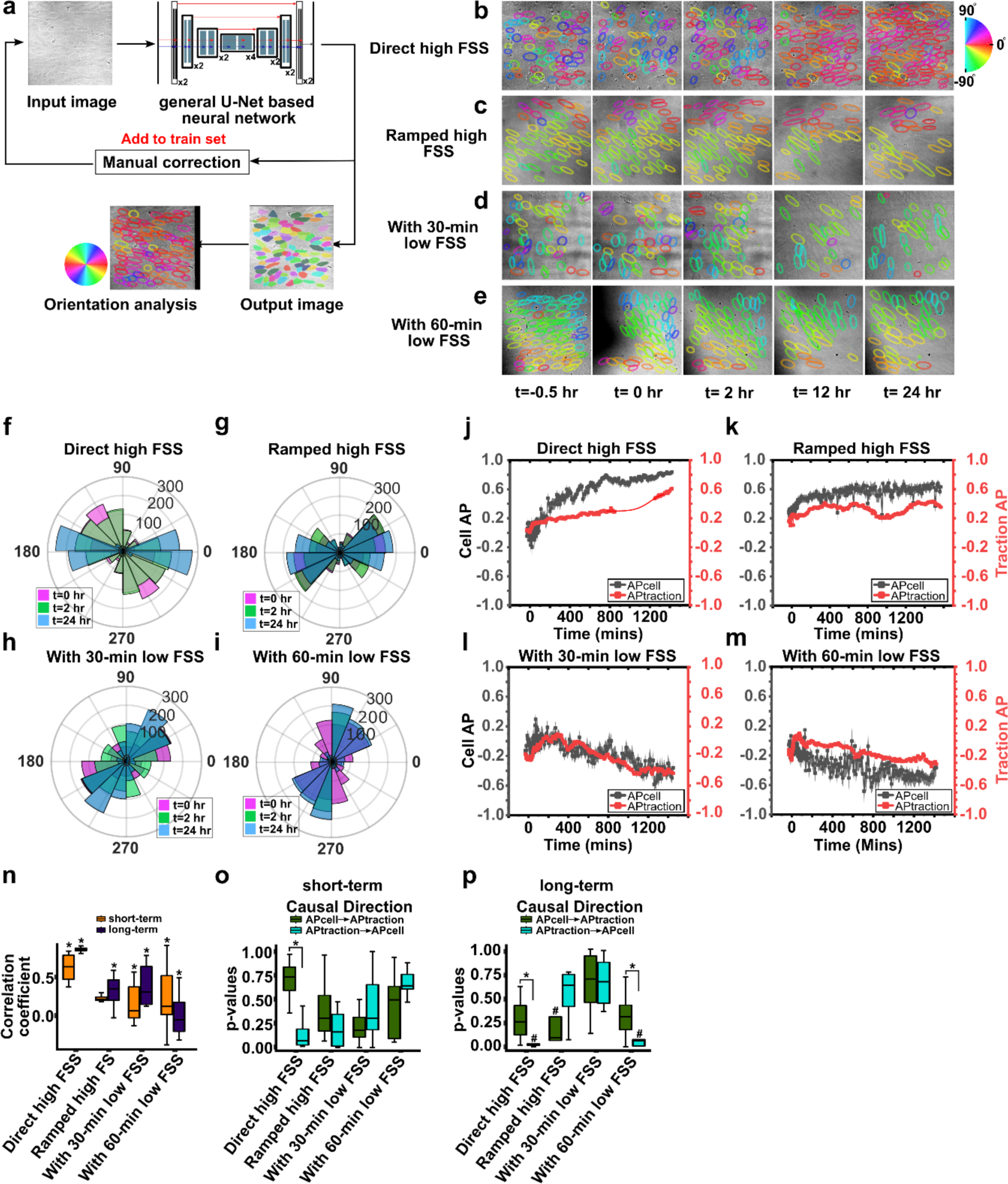
Traction alignment functionally causes cell alignment of HUVECs under direct high FSS or long-term 60-min low FSS. (a) Our cell orientation analysis framework consisting of Cellpose 2.0 neural network that uses a standard U-net backbone, which down-sample and upsample the feature maps along with skip connection between layers, with a human-in-loop procedure for manual training and correction. The output image after segmentation prediction was analyzed for orientation in custom developed python code for alignment parameter with angular distribution of the segmentation. (b-e) Predicted segmentation and ellipse fitting for direct high FSS (b), high FSS with ramp (c), 30-min low FSS (d), 60-min low FSS (e), conditions with angular color map showing the segmented cells with respective angular distribution color coding in the field of view for time t= -0.5 hr, 0 hr, 2 hr, 12 hr, 24 hr. (f-i) Polar histograms of cell orientations at t=0 hr (pink), at t=2 hr (green) and at t=24 hr (blue) for each condition with respect to flow direction. (j-m) Time-series plot of AP_cell_ (gray) and AP_traction_ (red) as a function of time from each representative time-lapse images recorded for direct high FSS (j, AP_traction_), High FSS with ramp (k, AP_traction_), 30-min low FSS (l, AP_traction_), and 60-min low FSS (m, AP_traction_). AP_traction_ for missing timepoints in panel j was calculated using non-linear curve fitting by gaussian amp function-R^2^=0.98404. Error bars: S.E.M. (n) A boxplot of Pearson correlation coefficients for short-term (orange) and long-term data (purple) for four FSS conditions. *: p < 0.05 tested with HMP-combined p-values of a robust test for zero-correlation^60^. (o-p) Boxplots of p-values from Granger Causality tests for short-term and long-term imaging for all conditions with causal direction that ‘AP_cell_ GCs AP_traction_’ (green) and ‘AP_traction_ GCs AP_cell_’ (blue). #: p < 0.05 tested with HMP-combined GC p-values, *: p < 0.05 tested with Mann-Whitney U-test between the two-direction GC test p-values. Sample sizes for each condition are N=4, 4, 6, 5, for direct high FSS, High FSS with ramp 30-min low FSS, 60-min low FSS, respectively.

The Granger Causality (GC)^62^ test was performed to determine whether AP_traction_ functionally causes AP_cell_ or vice versa. For our purpose, we analyzed the causality in both directions, *i.e.*, AP_cell_ granger-causes (GCs) AP_traction_, and, AP_traction_ GCs AP_cell_, from which a significance of the causality, *i.e.*, p-values, were produced. Again, in each experiment condition, the p-values were combined using the HMP to determine the global representative significance of each causal direction. A global representative significance less than 0.05 was considered significant. Furthermore, we checked if the two sets of GC p-values, representing the two GC directions, were significantly different. Since distributions of p-values were far from being Gaussian, the usual two-sample t-test could not be used. We instead employed the Mann-Whitney test^63^, which does not require parametric assumptions such as normality. These p-values are further combined using HMP. Combined p-values less than 0.05 are considered significant (Fig. 4o, p).

## Results

### Low FSS increases traction in the short-term but impairs force alignment

To investigate the fine-scale modulation of EC traction in response to changes in FSS, we developed a closed-loop flow system featuring a flow channel housing a silicone gel coated with fluorescent beads (Fig. 1a, Materials and Methods). We subjected a monolayer of HUVECs on this gel to varying FSS conditions and closely monitored their traction dynamics over time.

Initially, we sought to address conflicting findings in the literature regarding EC traction responses to high FSS. We applied a direct high FSS of 1 Pa to a HUVEC monolayer and observed traction modulation over a 90-minute period following the onset of flow. Differential interference contrast (DIC) images revealed that the monolayer did not exhibit significant gap formation between cells; instead, HUVECs continued to migrate while maintaining cell-cell boundaries after flow initiation (Fig. 1b, Video S1). Analysis of traction data showed a noticeable increase in traction magnitude, with traction vectors oriented both parallel and opposite to the flow direction (Fig. 1c, Video S2). Indeed, the average traction magnitude increased immediately after the onset of high FSS, plateauing approximately 30 minutes into the flow (Fig. 1d). This observed traction increase is consistent with previous reports^25,27,39,40^, albeit some of these findings were derived from long-term (≥20 hour) flow experiments^25,27^.

The abrupt application of high FSS, a common practice in many parallel-plate chamber studies^15,16,40,49^, could potentially lead to the detachment of ECs from neighboring cells or the extracellular matrix, affecting the transmission of traction^30,31^. To test whether this practice induces any difference in traction, we introduced a ramped FSS approach, gradually increasing FSS from 0 to 1 Pa within the first 30 minutes (Fig. 1e, *red profile*). HUVECs exposed to this ramped high FSS exhibited an immediate increase in traction, with the average traction plateauing within 20 minutes, even as FSS continued to rise (Fig. 1e, *black profile*). Furthermore, the average traction gain at the plateau was approximately half that observed with direct high FSS application (compare Fig. 1d *vs.* e). It is worth noting that differences in the initial, baseline traction between flow conditions, are within a normal fluctuation of the traction of HUVECs in a long-term static culture (Fig. S1). So the traction starting point was vertically translated at t=zero stating it as average traction. Similar trends, i.e., short-term rise in traction upon direct or ramped high FSS, were observed in two more replicate experiments per flow condition with 5 different viewing areas per experiment (Figs. S2 and S3). Thus, whether applied suddenly or in a ramped manner, high FSS induces a short-term increase in traction.

From the result from the ramped high FSS, *i.e.*, immediate traction elevation during the ramp phase where FSS is well-below 1 Pa, we hypothesized that such immediate rise in traction could be still demonstrated by HUVECs when facing low FSS. To test this hypothesis, we added an additional low-shear ramp for 30 min (Fig. 1f, *red*) or low-shear ramp followed by 30 min of steady low shear, totaling 60-min low shear, (Fig. 1g, *red*) before the high-shear ramp. As anticipated, both the 30-minute and 60-minute low FSS conditions showed a significant increase in traction during the initial low-shear ramp (Fig. 1f and g, *black*). The magnitude of the traction rises during the initial low shear ramp (5.5 Pa – 7.5 Pa in Fig. 1f,g) were comparable with the rise in the high FSS ramp case (∼7.5 Pa, from Fig. 1e), albeit smaller than the rise triggered by direct high FSS (∼16 Pa, Fig. 1d). Similar trends, i.e., intermediate short-term rise in traction upon low FSS were observed in two or more replicate experiments (Figs. S4 and S5). These results highlight that the introduction of low FSS elicits a substantial increase in traction as much as the amount observed in the ramped high FSS case.

Following these initial traction rises, HUVECs reached a traction plateau during the subsequent ramp to high FSS, a plateau that occurred at lower FSS levels for the 60-minute low FSS condition compared to the 30-minute low FSS condition (Fig. 1f,g, *blue lines and text*). Quantitative analysis of the FSS corresponding to maximum traction (termed FSS_@Tmax_, Fig. 1d-g, *blue lines and text*) revealed a decreasing trend from direct high FSS (FSS_@Tmax_ = 1 Pa) to ramped high FSS (FSS_@Tmax_ = 0.75 Pa) and further to 30-minute (FSS_@Tmax_ = 0.59 Pa) and 60-minute low FSS conditions (FSS_@Tmax_ = 0.24 Pa) (Fig. 1 m). This observation suggests that exposure to the lower FSS initially primes HUVECs for a reduced increase in traction when transitioning to high FSS conditions.

Cell and cytoskeletal alignment with the shear flow direction is a defining characteristic of ECs responding to high FSS^16,27,49,64^. Recent evidence has indicated that the alignment of traction vectors with the flow direction precedes cell shape alignment^45^, prompting us to consider traction alignment as an early marker for cell alignment. To analyze the traction field, we first examined the angular distribution of traction vector orientations. Prior to flow initiation, the distribution of traction vectors exhibited mostly isotropic characteristics (Fig. 1h-k). The application of direct high FSS induced a shift in the distribution towards the direction parallel to the flow within 90 minutes (Fig. 1h). Similarly, the high FSS condition with a 30-minute ramp showed an alignment shift towards the flow direction (Fig. 1i). However, with the introduction of low FSS for either 30 or 60 minutes, traction orientations shifted towards a more perpendicular alignment with respect to the flow (Fig. 1j, k).

To quantitatively assess the level of alignment, we introduced an alignment parameter, AP_traction_, for the traction field concerning a unit vector representing the flow direction (Materials and Methods). This parameter yields a value of 1 when all traction vectors align perfectly with the flow direction, -1 when they are perpendicular to the flow, and 0 when they exhibit an isotropic distribution. Analysis of AP_traction_ values per condition (Fig. 1n) revealed that direct high shear and the 30-minute ramp conditions exhibited a positive shift in alignment (Fig. 1h, i, o), whereas the introduction of low FSS resulted in either no significant alignment change or a negative shift in alignment, even after an additional 60 minutes under high FSS (Fig. 1j, k, o). Significant differences in change in AP_traction_ were observed between high FSS conditions and low FSS conditions towards the flow direction (Fig. 1o). These findings collectively suggest that exposure to low FSS impedes the traction remodeling process typically observed under high FSS, potentially resulting in cell misalignment.

### EC traction responses to FSS are shear-dependent and not nutrient-dependent

To ascertain that the observed changes in traction were indeed a direct consequence of FSS and not influenced by external factors such as nutrient availability, cytokine levels, oxygen concentration, or pH fluctuations, we conducted experiments involving the application of stepwise low FSS, ramped low FSS, or ramped high FSS for a duration of only 30 minutes, followed by flow cessation (Fig. 2a-c, *red*). The initiation of all three flow profiles elicited traction increases (Fig. 2a-c, *black, during first 30 minutes*) akin to the traction patterns observed in previous flow profiles culminating in high FSS conditions (Fig. 1d-f). Subsequent to flow cessation, ECs promptly exhibited a decline in traction (Fig. 2a-c, *black, after 30 minutes*), consistent with earlier traction observations following high and low FSS exposure^26^. These short exposures of low and high FSSs did not lead to noticeable traction alignment toward the flow direction after stoppage of the flow or 30 minutes after being under a static condition (Fig. 2 d-f). Consequently, the AP_traction_ value did not show any significant difference (Fig. 2g). Overall, the results suggest that the modulation of EC traction in response to flow is predominantly dictated by FSS and is minimally influenced by other extraneous factors.

### Exposure to low FSS governs long-term, secondary traction rise and relaxation

HUVECs showed a transient traction rise in the short term when exposed to both high and low FSS. This short-term response can be characterized as a temporary, transient reaction and has been anticipated to be succeeded by a prolonged, immune-suppressive relaxation^47^. Yet, a long-term traction response in addition to the short-term response of the same EC monolayer has not been elucidated. To unravel the intricate relationship between short-term and long-term effects, we subjected ECs to continuous exposure to 1 Pa FSS for 24 hours following the initial short-term FSS exposure, measuring and analyzing traction dynamics accordingly (Fig. 3). As anticipated, in the case of flow initiated with direct high FSS, the average traction magnitude began to decline after reaching a peak at 66 minutes into the flow, steadily decreasing to levels considerably lower than the initial traction observed before flow initiation (Fig. 3a). After 24 hours of flow, the traction had diminished to only approximately 30% of the initial traction level (Fig. 3a). This decrease in traction over prolonged high FSS exposure aligns with findings by Steward *et. al.*^45^.

Conversely, for flow initiated with a ramped high FSS, a secondary rise in traction occurred after the initial peak (Fig. 3b, *an arrowhead at t=250 min*), reaching a peak value (∼12 Pa), higher than the initial peak (∼8 Pa), and subsequently began to decrease over time, ultimately falling below the initial traction level before flow initiation (Fig. 3b), although not to the same extent as observed in flow initiated with direct high FSS. Similarly, when the flow began with a ramped low FSS followed by high FSS, the secondary rise in traction, as in the one observed with a ramped high FSS initiation, was observed (Fig. 3c) but with a longer time to reach the global maximum (Fig. 3c, *arrowhead at t=280 minutes*). After reaching this secondary peak, traction decreased, accompanied by a tertiary small rise. The traction took a longer time to return to the baseline traction than the flow with ramped high-FSS (Fig. 3c, *a blue asterisk at t=932 min*).

Interestingly, in the case of flow initiated with 60 minutes of low FSS, the EC traction exhibited not only a secondary rise but also a substantial tertiary rise (Fig. 3d, *arrowhead at t=600 minutes*). Consequently, the timing of each global traction maximum progressively increased with the duration of exposure to low initial FSS (Fig. 3i). Furthermore, after the tertiary peak, the elevated traction required an extended period to return to baseline levels. Remarkably, even after 24 hours, the traction remained well above the baseline traction (Fig. 3d). Consequently, the traction relaxation time, defined as the time needed for the average traction to recover to 110% of the baseline traction after flow, displayed a progressive increase corresponding to longer exposure to low initial FSS (Fig. 3j). These trends in traction magnitude suggest that exposure to low FSS, despite initially inducing a smaller transient rise, ultimately results in a substantially longer-lasting effect on traction (M=3 independent experiments, N=16-20 different cell monolayer positions per flow condition).

An examination of the polar distributions of traction fields revealed distinct responses to various flow conditions. When the flow was initiated with direct high FSS, the traction vectors notably reoriented towards the direction of flow (Fig. 3e), aligning with previous observations^45^. Similarly, the flow beginning with a ramped high FSS induced a similar alignment of traction vectors (Fig. 3f). Although the change in vector orientations, quantified by AP_traction_, was somewhat smaller in the representative cases of flow initiated with a ramped high FSS compared to direct high FSS (Fig. 3f vs. e), both exhibited significant positive changes relative to baseline (Fig. 3k).

In contrast, when the flow commenced with long-term exposure to 30 minutes or 60 minutes of low FSS, the traction orientation exhibited a more pronounced alignment perpendicular to the flow direction (Fig. 3g,h), accompanied by a reduction in the AP_traction_. Notably, the change in AP_traction_ was significant compared to an isotropic distribution, and the difference between these two conditions was not statistically significant (Fig. 3k). These findings align with previous observations that ECs align perpendicular to the flow in response to continuous low FSS for 16 hours^21^. These data suggest that a short-term exposure to low shear stress may trigger cytoskeletal remodeling resulting in random traction orientations that represent atherogenic conditions. Furthermore, our data also suggest that a prolonged initial low FSS exposure, *i.e.*, 30 min to 60 min, may require a considerably longer duration for ECs to align in the flow direction and diminish their traction responses.

### Traction alignment functionally causes cell alignment under direct high FSS and 60-min low FSS

To interpret the traction orientation phenotypes in relation to established flow-dependent endothelial behaviors, we evaluated EC cell orientation, as alignment with the flow direction is a known endothelial response to high FSS^16,27,49,64-66^. For this purpose, we employed a U-Net-based neural network, Cellpose 2.0^58^ for cell segmentation from DIC images, followed by analyzing their orientation (Materials and Methods, Fig. 4a). The analysis of cell orientation both before and after applying different flow profiles revealed clear distinctions. Direct application of high FSS resulted in significant alignment of cells along the flow direction over time (Fig. 4b,f). Ramped high FSS also led to cell alignment towards the flow, though the alignment was less pronounced (Fig. 4c,g). Conversely, when introducing low FSS for 30 and 60 minutes, there was a noticeable shift in alignment perpendicular to the flow direction (Fig. 4d,e,h,i). Notably, these patterns of cell alignment were similar to the observed distributions in traction orientation (Fig. 3e-f).

To determine the temporal relationship between cell vs. traction alignment, we analyzed the time-series of the APs of both cells and traction fields, AP_traction_ and AP_cell_, in a fine time scale. As expected, AP_cell_ increased over time under the conditions of the direct high FSS and the ramped high FSS with more distinct change observed in the direct high FSS (Fig. 4j,k). A similar trend was observed for AP_traction_: it increased over time in both direct high and ramped high FSS conditions, with larger increase for the direct-high FSS condition (Fig. 4j,k), which is consistent with the analysis in Fig. 3k. In contrary, both AP_traction_ and AP_cell_ decreased under the flow conditions started with low 30-min and 60-min FSS (Fig. 4l,m). The Pearson correlation between the pairs of the time series showed a strong positive correlation for the direct high FSS both for short-term and long-term (Fig. 4n). The conditions started with a ramped high FSS, 30-min low FSS and 60 min low FSS showed weaker but significant correlation except for short-term ramped high FSS (Fig. 4n), suggesting a tight coupling between traction alignment and cell alignment for nearly all flow conditions. A wide variance in the correlation coefficient for the short-term 30-min low FSS and both short- and long-term 60 min low FSS included some low coefficient values. However, the analysis with harmonic mean p-value (see Statistical Analysis section) yielded a significance.

To determine the functional causality between traction alignment and cell alignment, we performed Granger Causality (GC) test, a statistical hypothesis test used to determine if one time series can predict another^62^. This GC method, traditionally used in finance^67,68^ and neuroimaging^69^, has been adapted to a prediction in cell biology recently^70^. According to this test, if a time series X Granger-causes (GCs) Y, then changes in X will systematically occur before changes in Y. The test of each causal direction, *i.e.*, ‘AP_cell_ granger-causes (GCs) AP_traction_’, and, ‘AP_traction_ GCs AP_cell_’, produces a significance of causality as p-values. The p-values collected by several experiments per flow condition (Figs. S6-S9) were combined to determine the global representative significance of each causal direction with Harmonic Mean P (HMP)^61^. We calculated these p-values for AP time-series recorded in short-term (first 90, 90, 120 and 150 min for direct high FSS, ramped high FSS, with 30 min low FSS and 60 min low FSS after flow application, respectively, Fig. 4o) and in long-term (up to 24 hours, Fig. 4p). We separated the analyses because the two imaging modes had different time intervals, e.g., 1.5 min for short-term, and 10 min for long-term imaging. From the GC analysis with a short-term imaging, the combined p-value for the causal direction, AP_traction_ GCs AP_cell_, was not significant. However, the p-values were significantly smaller than p-values for the other direction (Fig. 4o). All other flow conditions in the short-term imaging did not show a significant difference between directions or in terms of the combined p-value. From the GC analysis with a long-term imaging, consistently, p-values for the causal direction, AP_traction_ GCs AP_cell_, were significantly lower than ones for the other causal direction, AP_cell_ GCs AP_traction_, (Fig. 4p), consistent with findings from the short-term response. Furthermore, the combined p-value for this direction, AP_traction_ GCs AP_cell_, was absolutely significant for direct high FSS (Fig. 4p). Similarly, in the 60 min low FSS condition, the combined p-value for the same direction, AP_traction_ GCs AP_cell_, was very significant. (Fig. 4p). The p-value of the Mann-Whitney test, testing the difference between the two sets of GC p-values in 60 min low FSS condition, was not significant but relatively small (0.1111). These results suggest that as long as the alignment becomes strongly along the flow (in case of direct high FSS) or perpendicular to the flow (in case of 60 min low FSS), traction alignment functionally causes cell alignment in a manner that is protective from, or prone to, the atherosclerosis. Taken together, these data suggest that traction alignment and cell alignment have a strong correlation and traction alignment is functionally responsible for cell alignment of HUVECs under direct high or 60-min low FSS introduction.

## Discussion

Utilizing a versatile flow system capable of tracking traction dynamics across multiple time scales, we have uncovered intriguing insights into the responses of ECs to varying FSS conditions. Our findings reveal that transient exposure to low FSS triggers an immediate short-term increase in traction, followed by a sustained traction rise when subjected to high FSS. In stark contrast, the application of direct or ramped high FSS initially provokes an increase in traction but subsequently leads to a marked decline in traction over the long term. Moreover, we observed that initial exposure to low FSS induces a distinctive traction field orientation, aligning more perpendicular to the flow direction, persisting even after the subsequent return to high FSS conditions. In contrast, direct or ramped high FSS prompts traction alignment parallel to the flow direction.

The short-term traction response to low FSS aligns with previous research that has demonstrated an immediate surge and subsequent decrease in traction upon the onset and cessation of low FSS, respectively^26^. This immediate traction response is believed to arise from the activation of FSS-triggered mechanotransduction and inflammatory signaling pathways. Notably, low FSS is known to canonically activate pathways such as NF-kB^71^, Smad2/3^18^ and the inflammatory mediator TNF-α^17^ while simultaneously reducing endothelial nitric oxide synthase (eNOS) production^72^ and ArhGap18^48^. A previous study has reported an increased traction with the introduction of TNF-α under 1.5 Pa of FSS^42^. Additionally, signaling mediated by Notch1, a well-established shear mechanosensor^73^, triggers RhoA signaling, subsequently activating myosin light chain and, consequently, myosin contractility and traction^40,74^. However, it is worth noting that some conventional low FSS-triggered transcriptional responses do not appear to be the direct cause of elevated traction but rather a consequence of it. For instance, research involving lipopolysaccharide (LPS)-based stimulation has shown that RhoA signaling operates upstream of NF-kB phosphorylation^75^. Furthermore, the Yap/Taz pathway, activated under low, oscillatory FSS^76^ via α5β1 integrin and tyrosine protein kinase ABL1 activation^77^, is known to be regulated by RhoA signaling^78^, potentially through cytoskeletal tension-mediated nuclear pore complex opening, facilitating Yap nuclear translocation^79^. Consequently, the rise in traction induced by low FSS may be attributed to molecular sensors activating pathways like Notch1 or TNF-α, which, in turn, activate RhoA, ultimately influencing NF-kB or Yap signaling. Further experimental investigations will be essential to unravel the intricate molecular circuitry underlying the sensing of low FSS by endothelial cells.

The occurrence of secondary or tertiary traction peaks observed predominantly in HUVECs subjected to low FSS pre-treatment bears a resemblance to the secondary peak documented in the activation of JNK2 signaling under high FSS conditions^24^. The JNK signal pathway operates downstream of TNF-α and upstream of key players such as c-JUN and NF-kB^80^. Notably, JNK activation is also modulated by factors downstream of integrin^24^ and RhoA activation^81^. Thus, the JNK2’s secondary peaks could be outcome of the secondary force peaks. Consequently, the secondary peaks observed in JNK2 activation may be a consequential response to secondary force peaks induced by high FSS. However, the precise upstream regulators responsible for generating these traction peaks remain elusive. Moreover, it remains unclear whether JNK2 exhibits similar secondary peaks in response to low FSS, and intriguingly, why such secondary peaks were not observed in our high FSS scenario. Our data reveals that the initial traction increase in response to low FSS is significantly less than half of the initial rise observed under direct high FSS conditions. This discrepancy leads us to speculate that the initial traction surge could potentially serve as a crucial signal triggering the initiation of an extensive remodeling program within the endothelial cells. Further investigations are warranted to unravel the intricate mechanisms governing these traction peaks and their potential roles in mechanotransduction processes.

The short-term initial exposure to low FSS has shown perpendicular alignment of the traction and cell to the flow direction even after 24 additional hours of high FSS. These results partially align with the study that shows perpendicular alignment of actin filaments to long-term low-FSS, disturbed flow^23,82,83^ where perpendicularly aligned ECs express high proinflammatory NF-kB^49^. Given the tight relationship between traction and cell alignment^45^, our results suggest that cytoskeletal remodeling initiated by gene reprogramming might have already begun with a short time exposure to low FSS. One such gene reprogramming in response to short-term low FSS could involve endothelial-to-mesenchymal transition (EndMT)^84-86^. EndMT leads to physiological changes like EC polarity and adherens junction (AJ) disruption. It will be worth investigating the possibility of EndMT, *e.g.*, by monitoring a EndMT-representing transcriptional factor, SNAIL^87^, with a range of time duration with low FSS before high FSS.

Reorientation of HUVEC traction and cell perpendicular to the flow after temporary low FSS exposure could be attempted to be explained by the set point theory^21,88,89^. The theory states that ECs have a preferred level of FSS and any deviation from the set point is compensated by vasoconstriction and dilation to maintain the set level of FSS^90^. According to this theory, it is possible that HUVECS set up their set-point FSS to be very low during the initial exposure to low FSS. The 1-Pa high FSS that is followed after the low FSS would be recognized as a too much FSS, more than a physiologically healthy level, which could result in EC alignment perpendicular to the flow, which has been shown to occur ECs under very high FSS^21^. Further experiments are needed to evaluate this interpretation.

In the context of ECs’ response to high laminar fluid shear stress (FSS), our findings reveal a dual-phase pattern: an initial short-term increase in traction followed by a long-term decrease. It’s important to note that these results do not serve to definitively reconcile the previously contradictory findings in the literature regarding endothelial traction responses^15,25,39,40,45^. Instead, they align with the established understanding of ECs’ shear-dependent behavior, wherein an initial immune response is succeeded by a long-term relaxation phase^47^. The variability in the current and previous shear-TFM results may be attributed to differences in the selection of cell types, flow conditions, ECM proteins, and the stiffness of the substrate. For instance, an BAEC monolayer on FN-coated surface has shown an increase in traction under 1.2 Pa FSS^25^ whereas on collagen-coated surface, HUVECs have shown a decrease in traction in a long-term^45^. Indeed, ECs on collagen-coated substrate have shown attenuated focal adhesion growth and cytoskeletal dynamics compared to ECs on fibronectin^91^. Moreover, ECs on collagen have shown transient activation of RhoA as compared to prolonged RhoA activation on fibronectin^92^. Thus, while the reasons for the inconsistent traction results is still undecisive, this ECM-dependence suggests that our finding of initial short-term traction rise is attributed to fibronectin’s synergistic integrin-activating function and associated RhoA activation, but it can still trigger, or be followed by, a long-term relaxation. It is also worth mentioning that all previous methods have measured traction at discrete time-points with at least one-minute interval, making it difficult to compare the remodeling response for the duration of FSS exposure whereas our results are from imaging done in a fine time-scale.

In conclusion, our data demonstrate unexpected shear-dependent traction modulation that temporary exposure to low FSS has a long-term effect featured by secondary rise in traction and reorientation of traction field perpendicular to the flow. While leaving many potential mechanistic possibilities in interpretation, these data suggest that traction itself might act as an important signal for endothelial pathophysiology in response to low shear stress.

## Acknowledgment

We thank Dr. Feng Zhao (Texas A&M University) for sharing early-passage HUVECs and helping us with HUVECs culture protocols. We thank M. A. Schwartz (Yale University) and Paul Evans (Imperial College London, UK) for their helpful discussion. We also thank Kathleen Pakenas in the lab for assisting the gel-fabrication process. This work was supported by NIH R15GM135806 grant. We also thank Health research institute (HRI) at Michigan Technological University for the graduate student fellowship.

## Supplemental Figures

**Figure S1.**
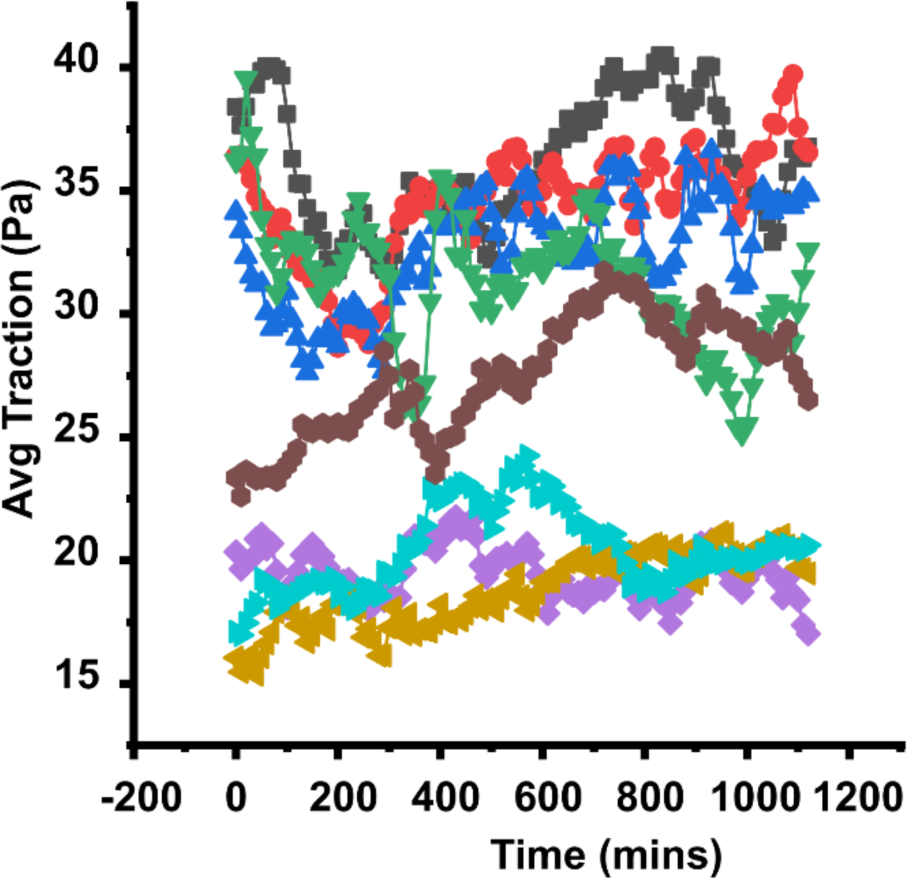
Traction fluctuation under long term static condition. Average traction magnitude generated by HUVECs monolayer under static condition with respect to time (N=2, M=8). Error bars: S.E.M. Here, N represents the number of flow experiments and M represents the number of regions of interest observed in each experiment.

**Figure S2.**
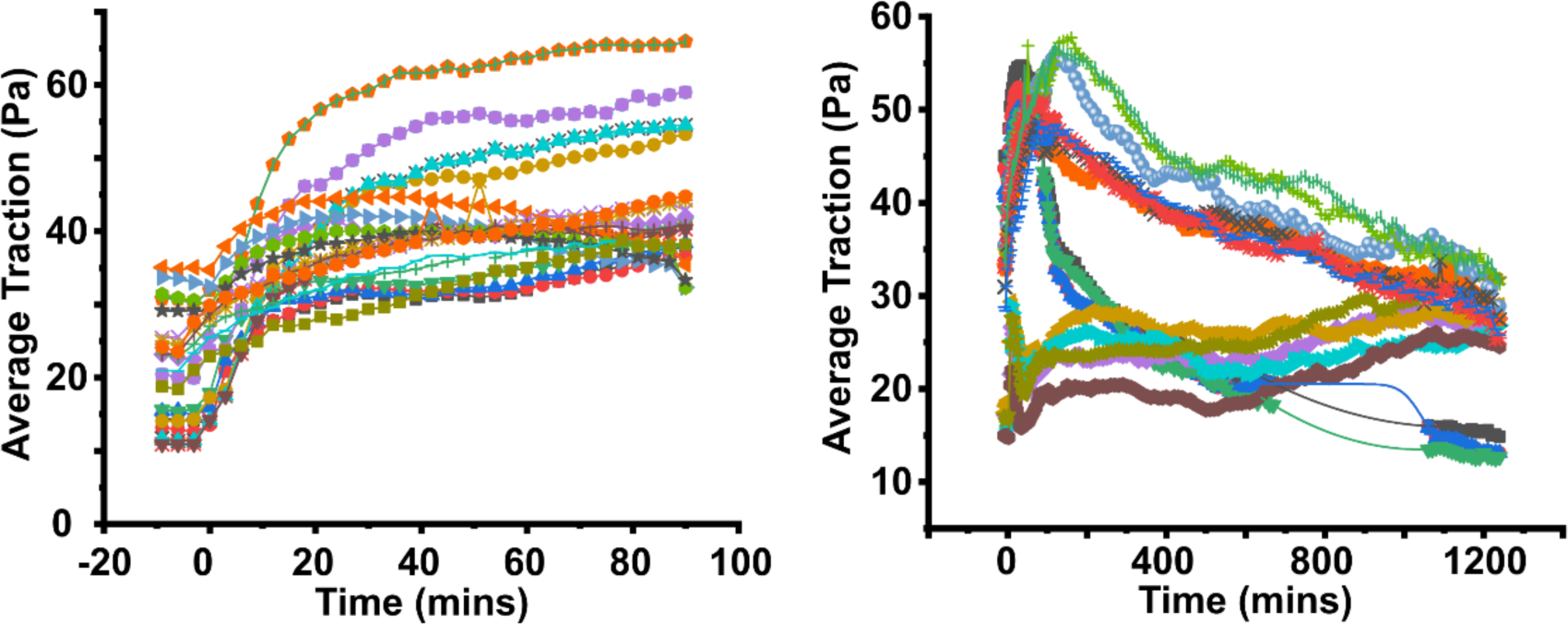
Average traction magnitude as a function of time for each individual region of interest, co-plotted with FSS profiles on ECs for direct 1 Pa FSS. (a) Short-term traction response (M=3, N=14). (b) Long-term traction response (M=2, N=9). Here, M represents the number of flow experiments and N represents the number of regions of interest observed in each experiment.

**Figure S3.**
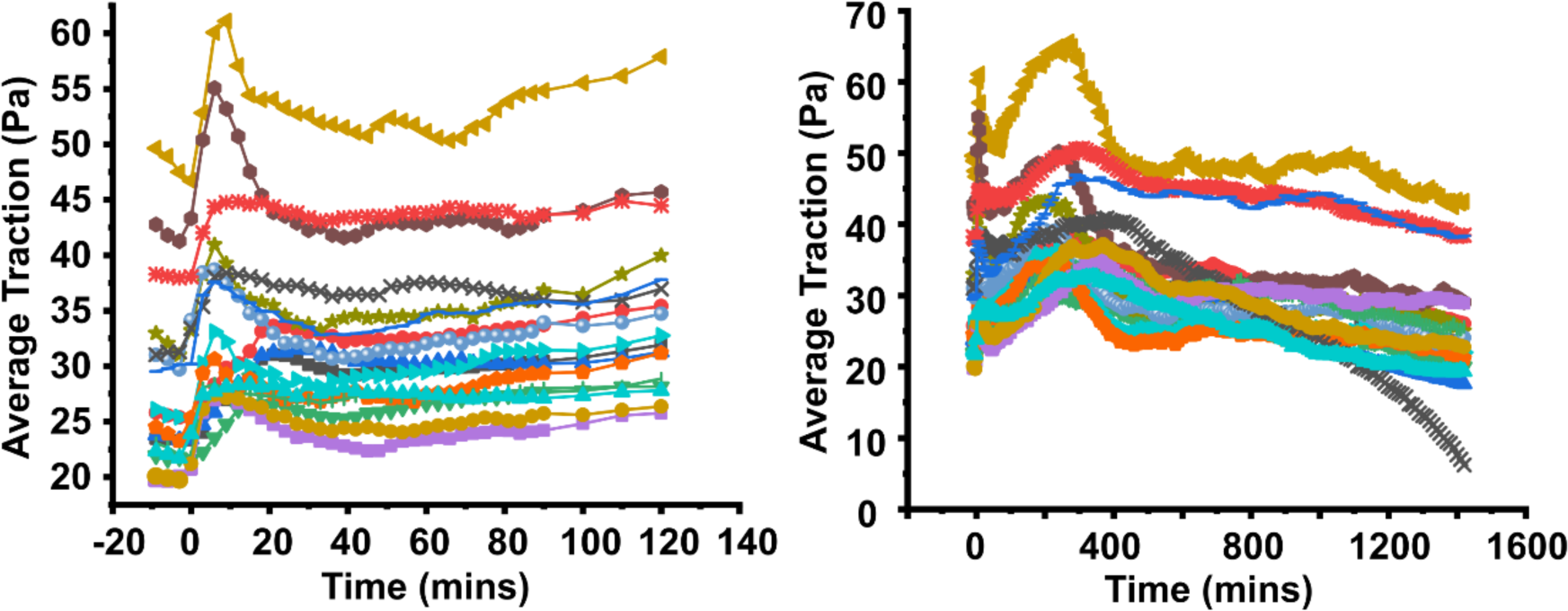
Average traction magnitude as a function of time for each individual region of interests, co-plotted with FSS profiles on ECs for 1 Pa FSS with 30 min ramp. (a) Short-term traction response (M=2, M=11). (b) Long-term traction response (M=2, N=11). M represents the number of flow experiments and N represents the number of regions of interest observed in each experiment.

**Figure S4.**
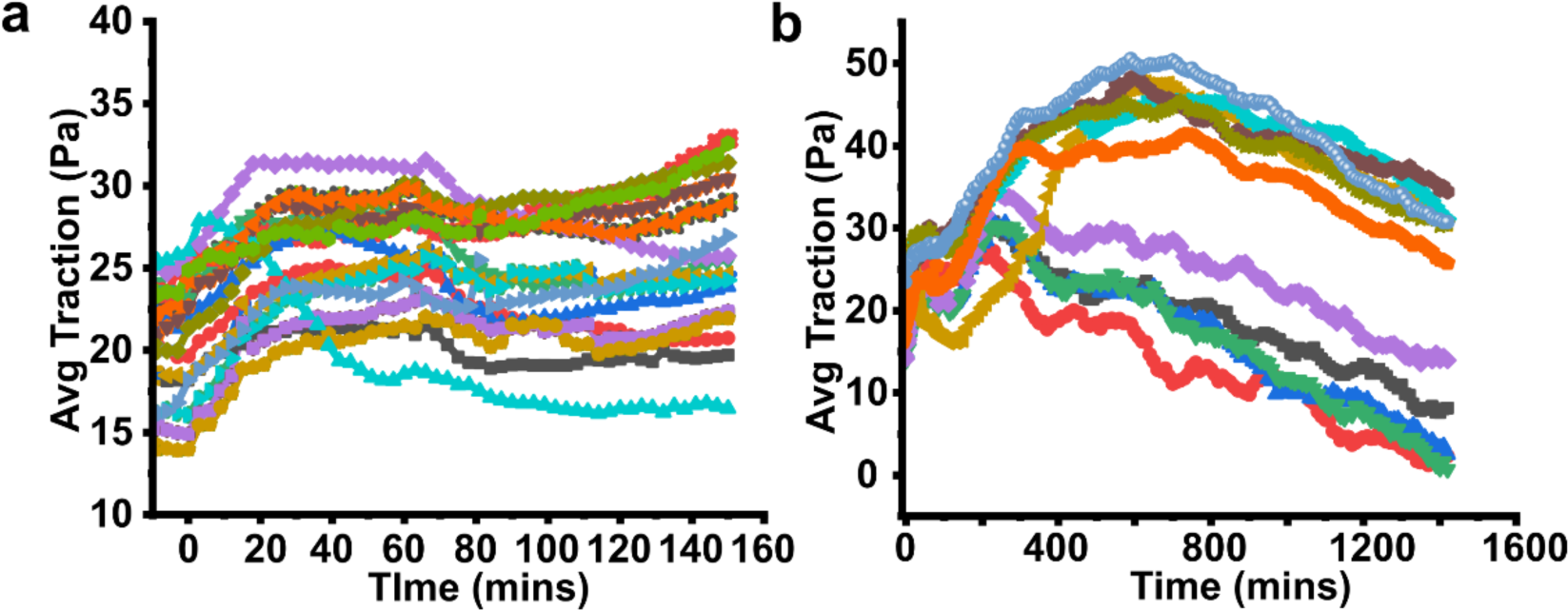
Average traction magnitude as a function of time for each individual region of interests, co-plotted with FSS profiles on ECs with 0.1 Pa FSS 30-min ramp. (a) Short-term traction response (M=4, N=38). (b) Long-term traction response (M=3, N=17). Here, M represents the number of flow experiments and N represents the number of regions of interest observed in each experiment.

**Figure S5.**
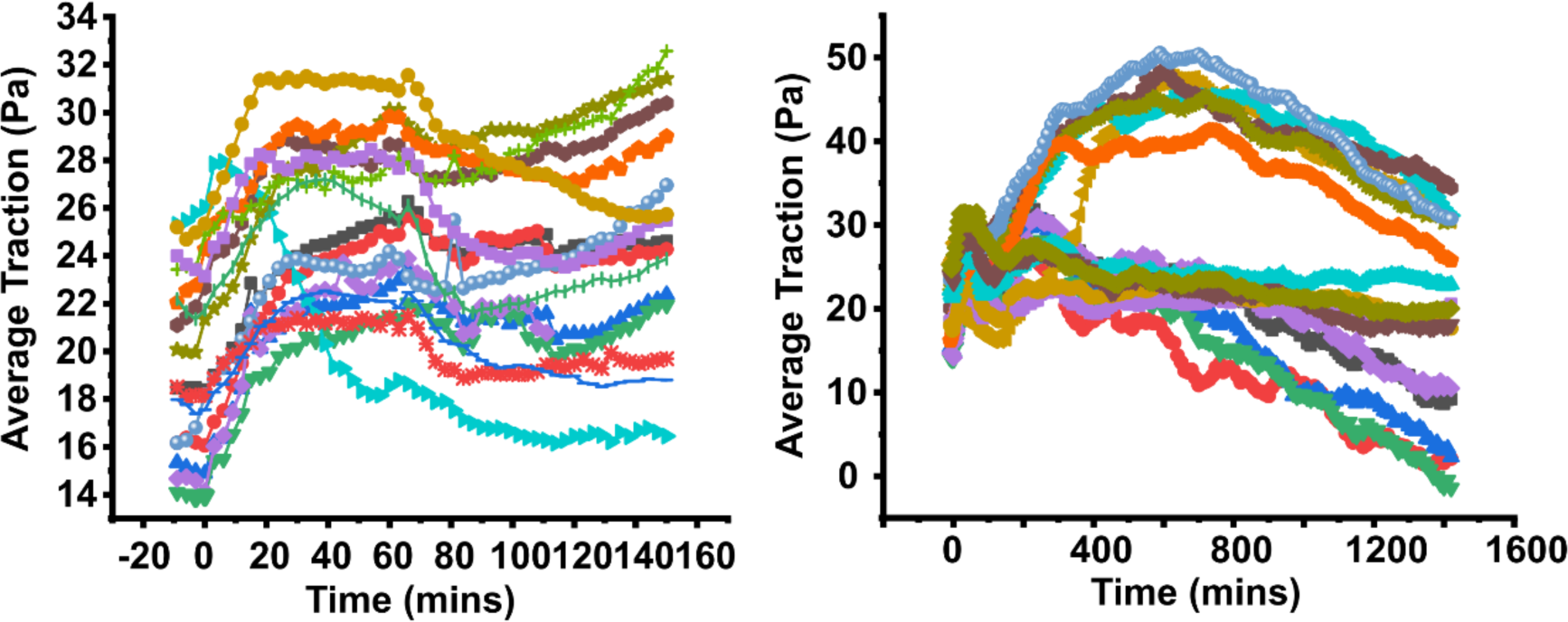
Average traction magnitude as a function of time for each individual region of interests, co-plotted with FSS profiles on ECs with 60-min 0.1 Pa shear. (a) Short-term traction response (M=4, N=24). (b) Long-term traction response (M=2, N=11). M represents the number of flow experiments and N represents the number of regions of interest observed in each experiment.

**Figure S6.**
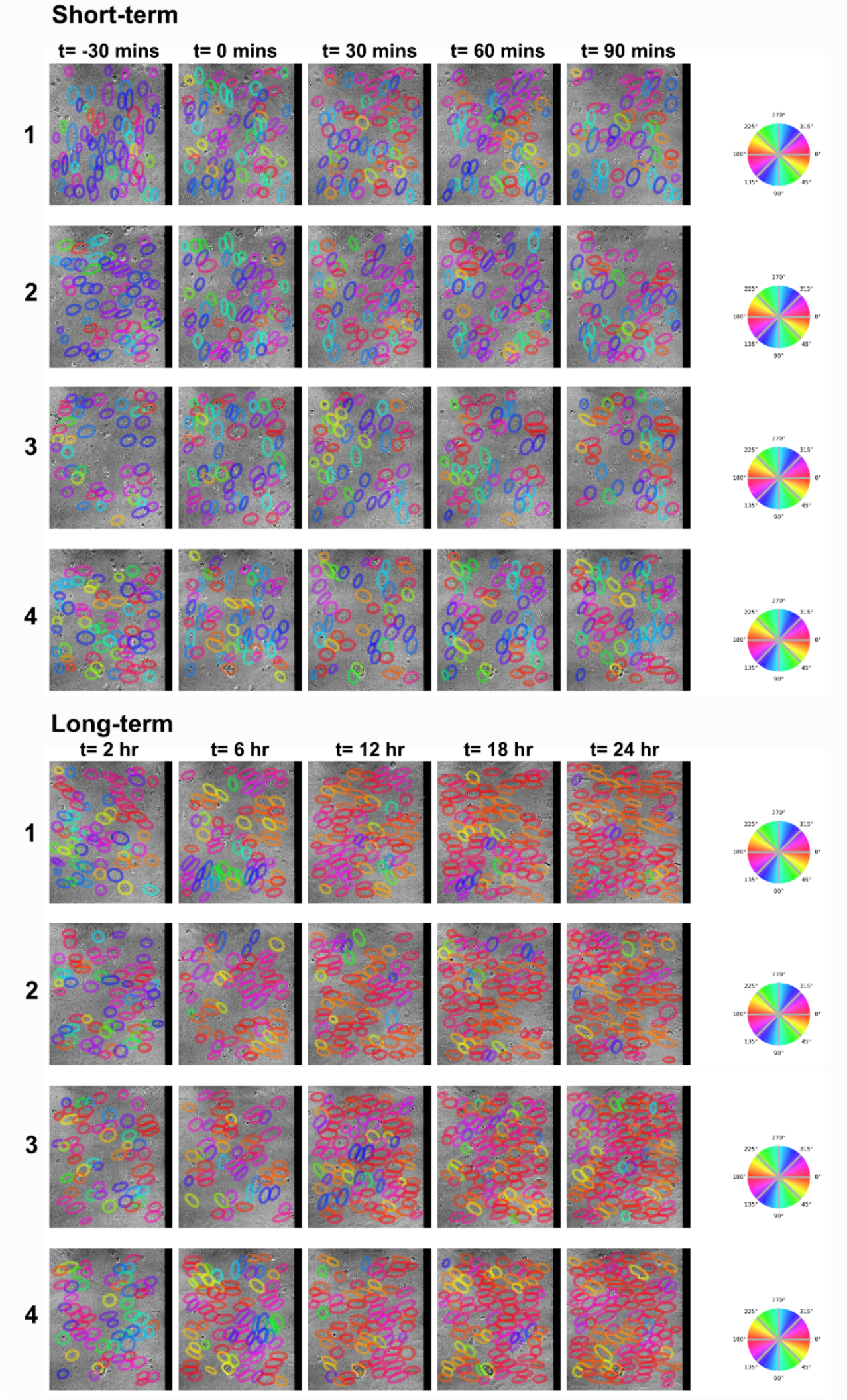
Predicted segmentation for direct high FSS for short-term and long-term with angular color map showing the segmented cells with respective angular distribution color coding in the field of view.

**Figure S7.**
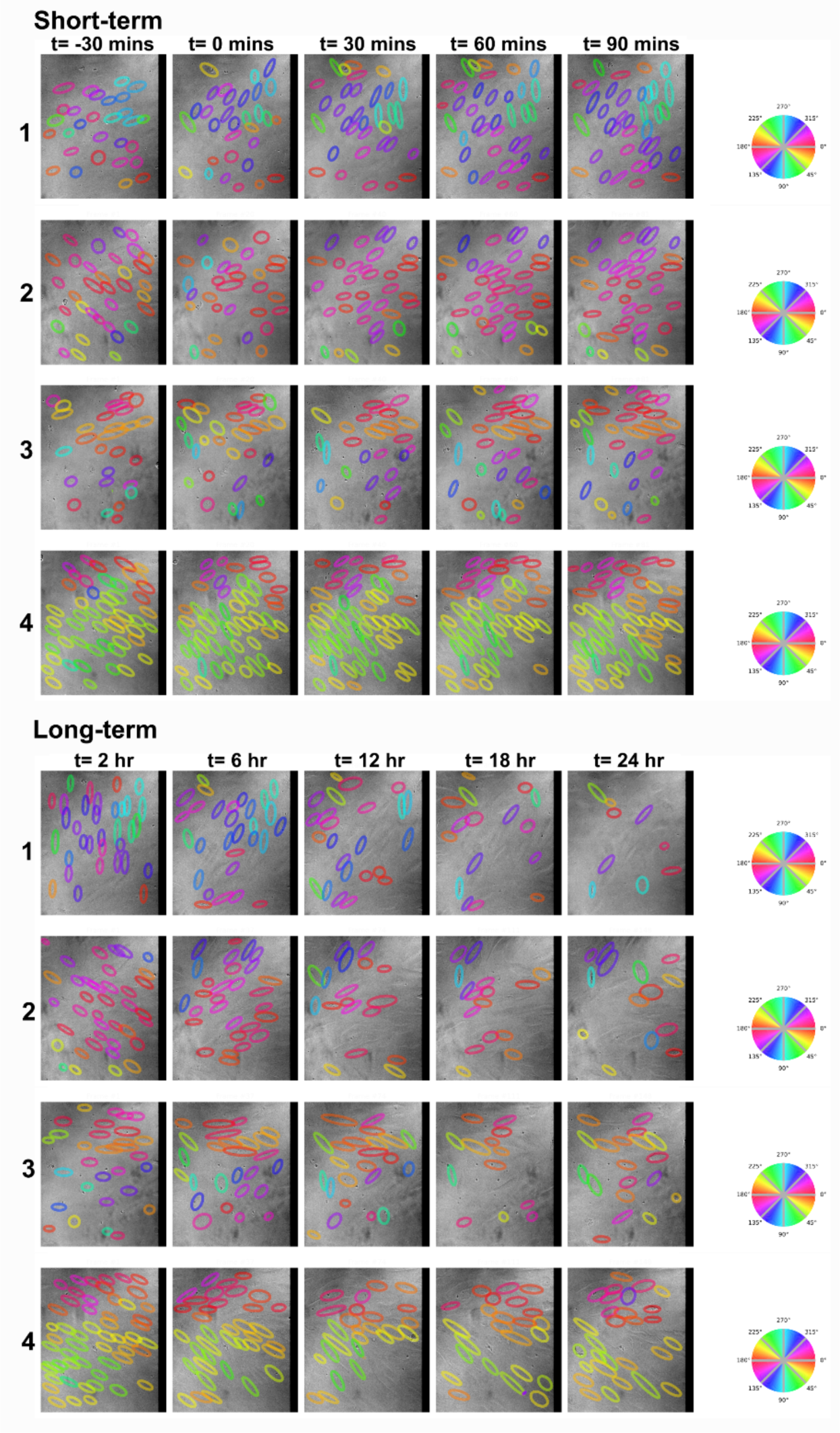
Predicted segmentation for high FSS with ramp for short-term and long-term with angular color map showing the segmented cells with respective angular distribution color coding in the field of view.

**Figure S8.**
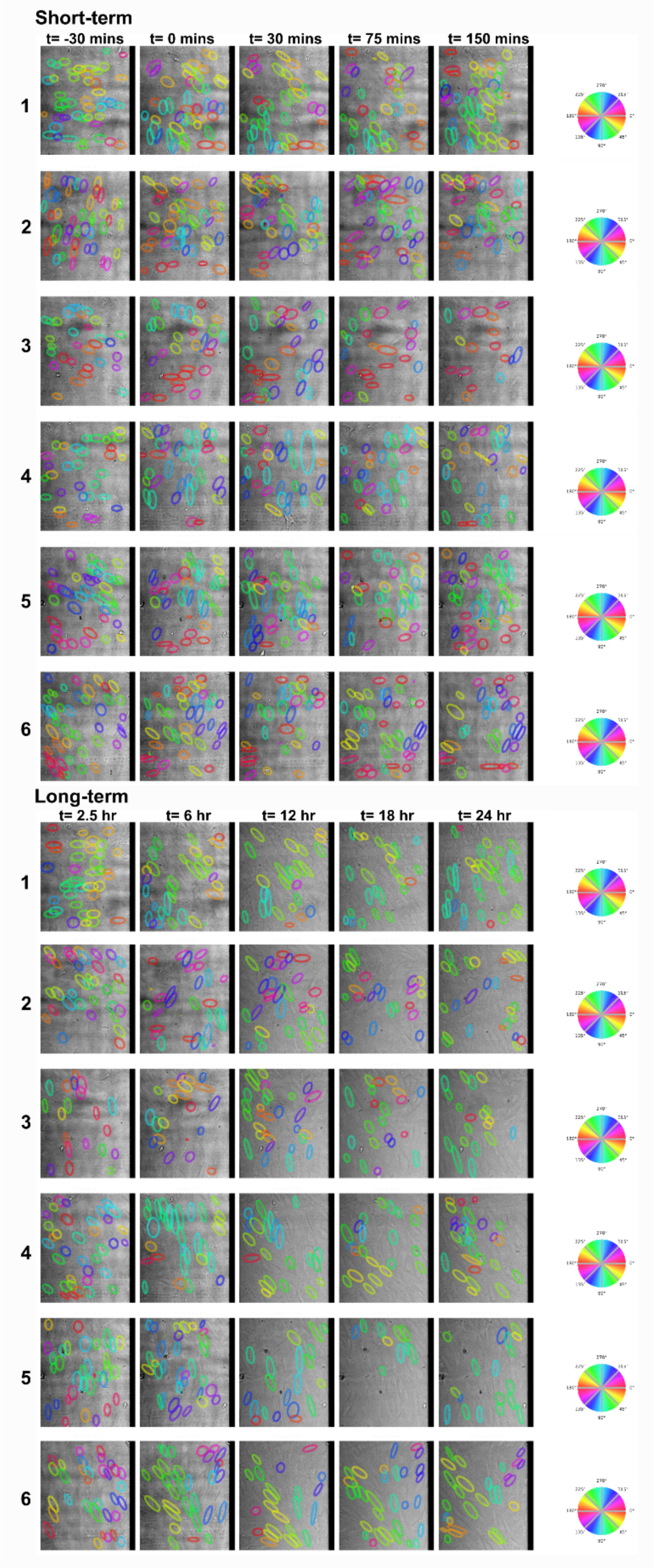
Predicted segmentation for 30-min low FSS for short-term and long-term with angular color map showing the segmented cells with respective angular distribution color coding in the field of view.

**Figure S9.**
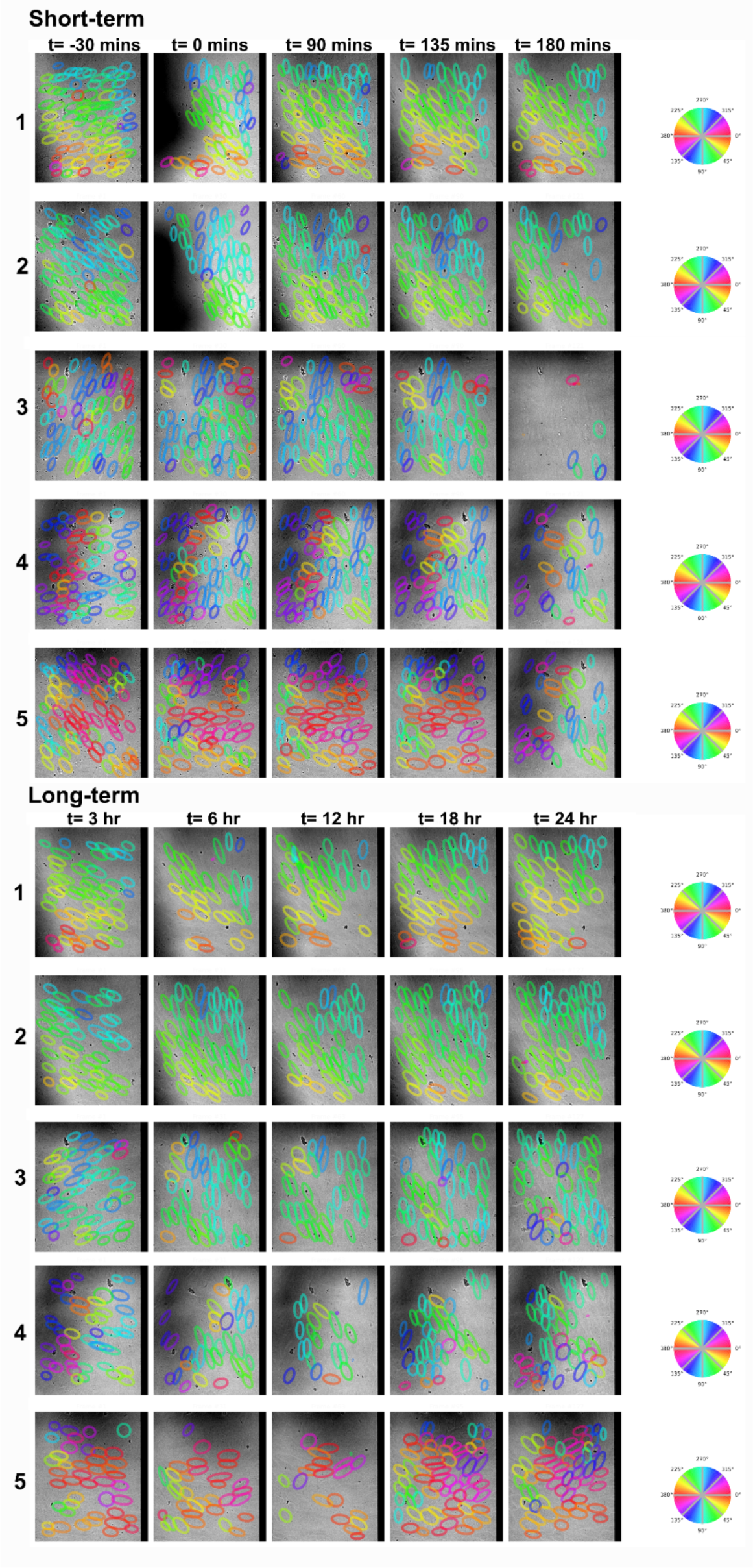
Predicted segmentation for 60-min low FSS for short-term and long-term with angular color map showing the segmented cells with respective angular distribution color coding in the field of view.

## Supplemental Video Legends

**Video S1.** Time-lapse DIC images of a region of a HUVECs monolayer on 2.6 kPa, bead-coated gel under direct high FSS. The time interval between consecutive frames: 90 seconds. Playing speed: 7 frames/sec. Total movie play time: 11 seconds. Duration of the movie: 90 minutes. Scale bar: 50 μm.

**Video S2.** Time-lapse color-coded traction vector field overlaid on DIC images of a region of a HUVECs monolayer on 2.6 kPa gel under direct high FSS. Time interval between consecutive frames: 90 seconds. Playing speed: 7 frames/sec. Total movie play time: 11 seconds. Duration of the movie: 90 minutes. Scale bar: 50 μm.

